# No sex differences in oxygen uptake or extraction kinetics in the moderate or heavy exercise intensity domains

**DOI:** 10.1101/2023.06.26.546455

**Authors:** Maria Solleiro Pons, Lina Bernert, Emily Hume, Luke Hughes, Zander Williams, Mark Burnley, Paul Ansdell

## Abstract

The integrative response to exercise differs between sexes, with oxidative energy contribution purported as a potential mechanism. The present study investigated whether this difference was evident in the kinetics of oxygen uptake (V□O_2_) and extraction (HHb+Mb) during exercise.

Sixteen adults (8 males, 8 females, age: 27±5 years) completed three experimental visits. Incremental exercise testing was performed to obtain lactate threshold and V□O_2peak_. Subsequent visits involved three six-minute cycling bouts at 80% of lactate threshold and one 30-minute bout at a work rate 30% between the lactate threshold and power at V□O_2peak_. Pulmonary gas exchange and near-infrared spectroscopy of the vastus lateralis were used to continuously sample V□O_2_ and HHb+Mb, respectively. The phase II V□O_2_ kinetics were quantified using mono-exponential curves during moderate and heavy exercise. Slow component amplitudes were also quantified for the heavy intensity domain.

Relative V□O_2peak_ values were not different between sexes (*p*=0.111). Males achieved ∼30% greater power outputs (*p*=0.002). In the moderate and heavy intensity domains, the relative amplitude of the phase II transition was not different between sexes for V□O_2_ (∼24 and ∼40% V□O_2peak_, *p*≥0.179) and HHb+Mb (∼20 and ∼32% ischemia, *p*≥0.193). Similarly, there were no sex differences in the time constants for V□O_2_ (∼28 s, *p*≥0.385) or HHb+Mb (∼10s, *p*≥0.274). In the heavy intensity domain, neither V□O_2_ (*p*≥0.686) or HHb+Mb (*p*≥0.432) slow component amplitudes were different between sexes.

The oxidative response to moderate and heavy intensity exercise did not differ between males and females, suggesting similar dynamic responses of oxidative metabolism during intensity-matched exercise.

**New and Noteworthy:** This study demonstrated no sex differences in the oxidative response to moderate and heavy intensity cycling exercise. The change in oxygen uptake and deoxyhaemoglobin were modelled with mono-exponential curve fitting, which revealed no differences in the rate of oxidative energy provision between sexes. This provides insight into previously reported sex differences in the integrative response to exercise.

## Introduction

The transition from rest to exercise involves an integrated response from the pulmonary, cardiovascular, and muscular systems to rapidly increase the supply and utilisation of oxygen for oxidative adenosine triphosphate (ATP) provision (Poole & Jones, 2012). The speed at which this process can occur can be quantified using pulmonary oxygen uptake (V□O_2_) kinetics, and is thought to determine metabolic stability and exercise tolerance across the spectrum of athletic performance and disease (Burnley & Jones, 2007; Grassi *et al*., 2011). The V□O_2_ response can be broken down into three phases, beginning with the initial cardio-dynamic phase (phase I, 10-20 s) which represents an increased venous return via the muscle pump effect, as well as increased pulmonary blood flow (Grassi *et al*., 1996). Thereafter, increases in pulmonary V□O_2_ are considered to reflect increased muscle oxygen uptake in response to exercise (phase II), until the energy demand of exercise is met by oxidative phosphorylation and V□O_2_ reaches a steady state (Hughson *et al*., 2001). A steady state response is attainable quickly within the moderate intensity domain, whereas in either heavy or severe intensity domains, a further rise in V□O_2_ is observed before the steady state is attained (heavy) or V□O_2max_ is reached (severe), termed the slow component. This three-phase response is ubiquitous in exercising humans; however, the biological characteristics of the individual can influence the rates at which they occur (for review see Poole & Jones, 2012).

The time constant of phase II kinetics are considered to be a crucial determinant of the decrease in contractile function experienced by the exercising individual (Temesi *et al*., 2017; Goulding *et al*., 2021), and could explain previously observed sex differences in the integrative response to exercise (Ansdell *et al*., 2020b). As oxidative phosphorylation does not immediately meet the demand for ATP, substrate-level phosphorylation is required (Burnley & Jones, 2007). Intuitively, the rate at which oxidative metabolism can be upregulated at the onset of exercise is inversely linked with the accumulation of deleterious metabolites such as hydrogen ions [H^+^] and inorganic phosphate [P_i_], as well as the depletion of phosphocreatine [PCr] stores, which all interfere with excitation-contraction coupling (Allen *et al*., 2008). However, the relationship between the aforementioned metabolites and the upregulation of oxidative phosphorylation is multi-faceted, with the progressive change in the phosphate energy state (i.e., the [ATP]/[ADP][P_i_] balance) driving the rate at which mitochondrial respiration increases, whilst [H^+^] accumulation concurrently inhibits anaerobic glycolysis at the onset of exercise (Tschakovsky & Hughson, 1999; Korzeniewski & Rossiter, 2015). Accordingly, Temesi *et al*. (2017) demonstrated a positive correlation between the time constant (τV□O_2_) of the phase II response and the decrease in quadriceps potentiated twitch force. Similarly, elite endurance athletes demonstrate faster compared to untrained individuals when exercising at similar relative intensities.

A consistent finding in studies comparing males and females exercising at the same metabolic intensity is that females experience a lesser degree of contractile impairment of the knee-extensors (Ansdell *et al*., 2019; Ansdell *et al*., 2020a; Azevedo *et al*., 2021). Previously, this has been suggested to be a result of sex differences in skeletal muscle composition, whereby females consistently demonstrate a greater proportional area of type I fibres (Staron *et al*., 2000; Roepstorff *et al*., 2006). The consequences of this fibre type difference are multi-factorial; for-instance, it is well established that type I fibres are more fatigue-resistant (Schiaffino & Reggiani, 2011). Additionally, female *vastus lateralis* capillary density is ∼23% greater in females compared to males (Roepstorff *et al*., 2006), while females also demonstrate greater mitochondrial oxidative function and intrinsic respiratory rates than males of equivalent training status (Cardinale *et al*., 2018). One factor that remains unexplored is whether these physiological sex differences result in differences in the metabolic response to exercise. Conceivably, the superior aerobic phenotype of female skeletal muscle could imply that females might be able to meet the ATP demand of exercise through oxidative means faster than males. Despite this, *ex vivo* evidence suggests that female skeletal muscle fibres have a lower ADP sensitivity of mitochondrial respiration (Miotto *et al*., 2018), which could result in a slower rate of oxidative phosphorylation at the onset of exercise. Therefore, *in vivo* assessment of V□O_2_ kinetics would provide insight into the balance between these morphological and cellular sex differences.

The V□O_2_ slow component is underpinned by different mechanisms to the phase II kinetics and describes the increase in V□O_2_ during constant-load exercise. This increase in V□O_2_ implies an impairment of efficiency and is likely an amalgamation of several concurrent physiological changes. Within skeletal muscle, the accumulation of metabolites (e.g., [Pi]) and associated contractile dysfunction is linked with the loss of efficiency (Grassi *et al*., 2015). Of relevance here, is that female skeletal muscle has consistently been demonstrated to be more fatigue-resistant (Ansdell *et al*., 2019; Ansdell *et al*., 2020a), and shows lesser increases in the surface electromyogram in states of fatigue (Ansdell *et al*., 2017; Ansdell *et al*., 2019). Furthermore, it is well-established that females have a greater reliance on lipid metabolism during sustained exercise at similar relative work rates (Cano *et al*., 2022), which could be related to the more oxidative phenotype of skeletal muscle, as described above. Combined, these physiological sex differences could represent a slower loss of efficiency during constant load exercise in females, however, this remains unexplored.

Despite more aerobically-suited skeletal muscle, females have lower levels of haemoglobin (Murphy, 2014), which is thought to impair O_2_ carrying capacity during exercise (Harms *et al*., 1998; Diaz-Canestro *et al*., 2022). During exercise where O_2_ delivery and utilisation are both limiting factors (e.g., cycling, Goulding & Marwood, 2023), these factors are thought to counteract each other to enable comparable relative metabolic thresholds (i.e., critical power) between the sexes (Ansdell *et al*., 2020b). To date, the only investigation to systematically investigate the oxidative adjustment at the onset of exercise between sexes did so during low intensity treadmill walking (Beltrame *et al*., 2017). Data from this study suggested faster O_2_ extraction in females, quantified as the change in de-oxyhaemoglobin and myoglobin (HHb+Mb) signal in near-infrared spectroscopy (NIRS), fitting with the notion that phase II V□O_2_ kinetics are influenced by intramuscular factors in healthy humans (Poole & Jones, 2012). However, the demands of treadmill walking differ to those of high-intensity cycling exercise, where it has recently been argued that all levels of the O_2_ cascade are considered to be influential in determining metabolic responses to exercise (Goulding & Marwood, 2023). Thus, given the lack of evidence regarding physiological responses to exercise in females (James *et al*., 2023), it remains to be determined how sex differences in convective and diffusive contributions to O_2_ delivery mediate the rate of V□O_2_ adjustment to exercise.

Accordingly, the present study employed a multi-method approach of measuring pulmonary gas exchange and near-infrared spectroscopy simultaneously to compare the kinetics of pulmonary V□O_2_ as well as muscle oxygen extraction in both sexes during moderate and heavy intensity exercise. Previously this experimental approach has been used to obtain information about O_2_ delivery and utilisation to gain insights into the integrative response to exercise (DeLorey *et al*., 2003). It was hypothesised that females would demonstrate a smaller value for the phase II time constant (i.e., faster kinetics) for V□O_2_ and HHb+Mb at the onset of exercise, and a smaller slow component amplitude in the heavy intensity domain.

## Methods

### Ethical Approval

This study received institutional ethical approval from the Northumbria University Health and Life Sciences Research Ethics Committee (submission reference: 49189) and was conducted according to all aspects of the Declaration of Helsinki, apart from pre-registration in a database. Participants volunteered for the study and provided written informed consent.

### Participants

Using the effect size for the sex difference in *vastus lateralis* tissue oxygenation during heavy intensity exercise (ηp^²^ = 0.509) from (Ansdell *et al*., 2020a), an *a priori* sample size calculation determined a minimum of 14 participants (seven females and seven males) were required to detect an effect (α = 0.05, power = 0.95). Therefore, eight males (mean ± SD age: 27 ± 3 years; stature: 182 ± 5 cm; body mass: 75.3 ± 10.2 kg) and eight females (mean ± SD age: 27 ± 7; stature: 163 ± 4 cm; body mass: 61.8 ± 5.9 kg) volunteered to take part in the study. Hormonal status was not an exclusion criterion or controlled for in this study. Female participants were tested in any phase of their menstrual cycle and there were no restrictions on hormonal contraceptive usage. This decision was based on evidence from (Mattu *et al*., 2020) who demonstrated no hormonal effects on V□O_2_ kinetics during cycling exercise. Of the eight females, two were using combined oral contraceptive pills (Lucette and Rigevidon) and six were naturally menstruating. All participants were free from cardiovascular, respiratory, and neurological disease as well as musculoskeletal injury.

### Experimental Design

All participants visited the laboratory on three occasions across an average of 10 ± 4 days (range: 5 – 21 days). During the first visit, participants were familiarised with the experimental procedures and completed two incremental exercise tests to quantify lactate threshold and peak oxygen uptake (V□O_2peak_). The second and third visits were identical and involved three six-minute bouts of moderate intensity exercise (80% of lactate threshold, LT), separated by six minutes of unloaded pedalling. Thereafter, a single bout of heavy intensity exercise (30% ΔLT – V□O_2peak_) was performed for 30 minutes.

### Visit 1: Familiarisation & Incremental Testing

The first visit began with participants completing a screening questionnaire to ensure inclusion criteria were met. Thereafter, participants moved onto the cycle ergometer (Velotron, SRAM, Chicago, IL, USA) which was set up with the seat height aligned with the hip, and handlebar height set according to the participants’ comfort, these measurements were recorded and replicated for subsequent trials. The breath-by-breath gas exchange mask was then placed over the participant’s mouth and nose, and an air-tight seal was ensured before resting data was recorded. Following resting measures of pulmonary gas exchange and muscle oxygenation, participants completed five minutes of warm-up cycling at a light intensity (60 W) at a self-selected cadence between 70-100 rpm, before commencing an incremental exercise test. The first incremental exercise test began at 75 W and increased by 25 W every five minutes. At the end of each stage, a capillary blood sample was drawn from the participants’ fingertip and immediately analysed to determine whole blood lactate concentration (mmol.L^-1^, Biosen C-Line, EKF Diagnostics, Germany). The test was terminated once LT was identified as the first work rate at which a non-linear increase in blood lactate concentration was observed (Faude *et al*., 2009), after which, participants were provided 20 minutes of passive rest.

Next, participants began the second incremental test with five minutes of warm-up cycling at a light intensity (60 W) at the same self-selected cadence as before. Thereafter, a ramp test beginning at 75 W commenced, with power output increasing 1 W every 2.4 seconds (25 W.min^-1^). This test was terminated at volitional exhaustion, defined as cadence falling >10 rpm for five seconds. Strong verbal encouragement was provided to participants throughout. A final blood lactate sample was drawn immediately after volitional exertion. The greatest 30 second average V□O_2_ value was used to quantify V□O_2peak_, whilst the final power output was used to quantify maximal ramp test power (Pmax).

### Visits 2 & 3: Square-Wave Exercise Bouts

Visits two and three were identical, and performed with a minimum of 24 h between visits. The visits involved continuous sampling of pulmonary gas exchange and near infrared spectroscopy (NIRS) of the *vastus lateralis*. Trials commenced with participants performing three minutes of unloaded pedalling on the cycle ergometer. Thereafter, participants performed three repetitions of six-minute cycling bouts at 80% of the work rate associated with LT (moderate intensity exercise), interspersed with six minutes of unloaded pedalling. Following this, participants cycled for 30 minutes at a work rate 30% between the LT and V□O_2peak_ (30%Δ, heavy intensity exercise). Throughout this visit, participants were asked to replicate their self-selected cadence from visit 1, which was monitored by an experimenter throughout. Exercise intensity was altered abruptly in a ‘square-wave’ fashion for each repetition.

Immediately following all exercise, a ‘physiological calibration’ of NIRS signals was performed as per recommendations from Barstow (2019). Participants were laid supine on a physiotherapy table, with their leg placed horizontal, and an automatic personalised tourniquet system for blood flow restriction (Delfi Medical Innovations Inc., Vancouver BC, Canada) was placed around the thigh via a nylon cuff (11.5 cmC:×C:86 cm, 5 mm thick), proximal to the NIRS optode. The cuff was inflated for five minutes at 120% of limb occlusion pressure in order to occlude blood flow, and the mean pressure was not different between males and females (120% limb occlusion pressure: 248 ± 20 vs. 249 ± 30 mmHg, *p* = 0.780). This system automatically measures limb occlusion pressure, defined as the minimum pressure required for complete restriction of arterial blood flow in a limb, and maintains the pressure during inflation to ensure consistent occlusion. Following this, pressure in the cuff was released and the hyperaemic response measured (see *Near Infrared Spectroscopy*). This protocol allowed all NIRS data to be expressed as a % of an individual’s physiological minimum and maximum values. Physiological calibration negates any potential influence of adipose tissue thickness on NIRS data (Barstow, 2019).

### Pulmonary Gas Exchange

During all visits, expired gas was analysed breath-by-breath using an online system (Vyntus CPX, Jaeger, CareFusion, Germany). Oxygen (O_2_) and carbon dioxide (CO_2_) concentrations were quantified via a paramagnetic chemical fuel cell and non-dispersive infrared cell respectively. Before each test, the analysers were calibrated using ambient air and a gas of known O_2_ (15.00%) and CO_2_ (4.97%) concentrations. Ventilatory volumes were inferred from measurement of gas flow using a digital turbine transducer (volume 0 to 10 L, resolution 3 mL, flow 0 to 15 L·s^-1^) and calibrated prior to each test (Hans Rudolph Inc. Kansas City, USA).

### Near Infrared Spectroscopy

A multi-distance, continuous-wave, single channel NIRS (NIRO-200NX, Hamamatsu) was used to evaluate changes in vastus lateralis muscle deoxyhaemoglobin and myoglobin (HHb+Mb), as well as oxyhaemoglobin and myoglobin (HbO_2_+MbO_2_) concentrations, sampled at a rate of 5 Hz. Tissue oxygenation index (TOI) was calculated as HbO2+MbO2 ÷ [HbO2+MbO2 + HHb+Mb] × 100. The light-emitting probe comprised of light emitting diodes operating at three wavelengths (735, 810, and 850 nm). The probe was placed on the *vastus lateralis*, 20 cm above the fibular head. Optodes were held in place by an elasticised bandage and covered by an opaque, dark material to avoid motion and ambient light influences.

### Data Analysis

#### V□O_2_ Kinetics

The breath by breath data was manually filtered to remove outlying breaths, defined as breaths deviating more than 500 ml.min^-1^ from the mean value from the preceding five breaths. Thereafter, breath by breath data was linearly interpolated to provide second-by-second values. The multiple repetitions of the square-wave exercise bouts were then averaged, and V□O_2_ responses were time aligned to the onset of exercise. Data from the onset of the transition to 20 s was removed, then the resultant data was modelled with a monoexponential curve, including data from -60 to 360 seconds (moderate) or -60 to 120 seconds (heavy), with the following equation:

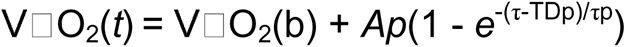

Where V□O_2_(t) is the V□O_2_ at time *t*; V□O_2_(b) is the baseline V□O_2_ measured in the 60 s preceding the transition in work rate; and *A_p_*, TD*_p_*, and τ_*p*_ are the amplitude, time delay, and the time constant of the phase II response, respectively. We chose to constrain the modelling of heavy intensity domain onset kinetics to 120 seconds in order to minimise the influence of the slow component, however this cannot be guaranteed (Burnley *et al*., 2006). For exercise in the heavy intensity domain, the amplitude of the V□O_2_ slow component was determined by subtracting the phase II amplitude from the highest 30 s average of V□O_2_ during the 30 min bout (Rossiter *et al*., 2001). To facilitate comparisons between sexes, amplitudes were also normalised to each individual’s V□O_2peak_ and end-exercise V□O_2_ as well as being presented in L.min^-1^. The O_2_ cost of the transitions was estimated by calculating the V□O_2_ gain (Porcelli *et al*., 2016), where the amplitude of the phase II response was divided by work rate (ml.min^-1^.W^-1^).

### Deoxyhaemoglobin Kinetics

Prior to curve fitting, data from NIRS were normalised to the minimum values during, and the maximum values following the five minute arterial occlusion (Ryan *et al*., 2012), then averaged into 1 s and 5 s bins. The multiple repetitions of the square-wave exercise bouts were then averaged, and HHb+Mb responses were time aligned to the onset of exercise. The TD for the HHb+Mb response was determined using the 1 s averaged data as the time between exercise onset and the first point at which HHb+Mb signal started to systematically increase. This was performed for each transition individually, with all TDs averaged to provide a single value. The 5 s averaged data was then modelled O_2_ data, including data up to 90 s after the transition (Murias *et al*., 2010). Other NIRS-derived variables (TOI and HbO_2_+MbO_2_) were quantified as 30 s averages at the following time points: the 30s of unloaded pedalling immediately prior to the transition, 90 s following the transition (heavy intensity domain only), and the final 30s of the square wave bout of exercise.

### Statistical Analysis

Data are presented as mean ± SD within the text and figures. Normal distribution of data was confirmed with the Shapiro-Wilk test. As all variables had normally distributed data, males and females were compared with independent samples t tests for variables with a single value or time point. For repeated measures variables during exercise, two-way (sex × time) repeated measures ANOVA were performed, followed by Bonferroni-corrected post hoc tests if significant main effects were observed. Effect sizes for comparisons were calculated as Cohen’s *d.* The significance level for all statistical tests was set at *p* < 0.05.

## Results

### Incremental Exercise Testing

Anthropometric data and outcome variables from the two incremental exercise tests performed in the first visit are presented in Table 1. As expected, males had a greater stature and body mass than females (*p* ≤ 0.006) as well as a greater absolute V□O_2peak_ (mean difference: 39%, *p* = 0.002). However, when V□O_2peak_ was expressed relative to body mass, no sex difference was observed (mean difference: 14%, *p* = 0.111). Males also exercised at greater power outputs than females, with Pmax and LT being ∼30% greater in males (*p* ≤ 0.023), however when LT was expressed as a % of Pmax, no sex difference was observed (*p* = 0.373). This resulted in the power outputs for the moderate intensity bouts being greater in males compared to females (*p* ≤ 0.023). and heavy

**Table 1:** Anthropometric data and outcome variables from incremental exercise testing.

|  | Males (n = 8) | Females (n = 8) | P value | Cohen's d |
| --- | --- | --- | --- | --- |
| Age (years) | 27 ± 3 | 27 ± 7 | 0.734 | 0.117 |
| Stature (cm) | 182 ± 5 | 163 ± 4 | **< 0.001** | **1.383** |
| Body Mass (kg) | 75.3 ± 10.2 | 61.8 ± 5.9 | **0.006** | **0.501** |
| $\dot{V}\text{O}_{2\text{peak}}$ (L.min<sup>-1</sup>) | 3.47 ± 0.58 | 2.50 ± 4.42 | **0.002** | **0.627** |
| Relative $\dot{V}\text{O}_{2\text{peak}}$ (ml.kg<sup>-1</sup>.min<sup>-1</sup>) | 46.2 ± 6.6 | 40.5 ± 6.7 | 0.111 | 0.326 |
| Pmax (W) | 328 ± 54 | 236 ± 43 | **0.002** | **0.646** |
| Pmax (W.kg<sup>-1</sup>) | 4.4 ± 0.8 | 3.8 ± 0.5 | 0.118 | 0.253 |
| Power at LT (W) | 153 ± 25 | 119 ± 29 | **0.023** | **0.528** |
| LT (% Pmax) | 47 ± 6 | 50 ± 7 | 0.373 | 0.183 |
| LT (W.kg<sup>-1</sup>) | 2.0 ± 0.2 | 1.9 ± 0.4 | 0.440 | 0.127 |
| 80% LT (W) | 123 ± 20 | 95 ± 23 | **0.023** | **0.528** |
| 80% LT (W.kg<sup>-1</sup>) | 1.6 ± 0.2 | 1.5 ± 0.3 | 0.440 | 0.110 |
| 30% Δ (W) | 206 ± 30 | 154 ± 32 | **0.005** | **0.653** |
| 30% Δ (W.kg<sup>-1</sup>) | 2.7 ± 0.4 | 2.5 ± 0.4 | 0.198 | 0.213 |

LT: lactate threshold, Pmax: Maximal ramp test power output, $\dot{V}\text{O}_{2\text{peak}}$ : Maximal rate of oxygen consumption.

### V□O_2_ Kinetics

The transition from unloaded pedalling to moderate and heavy intensity cycling elicited an increase in V□O_2_ (see Figure 2), and the monoexponential curve used to describe the increase in V□O_2_ in males and females demonstrated excellent r^2^ values (see Table 2).

**Table 2:** Data from the monoexponential modelling of V□O_2_ kinetics during moderate and heavy intensity transitions.

|  | Males (n = 8) | Females (n = 8) | P value | Cohen's d |
| --- | --- | --- | --- | --- |
| *Moderate Intensity Domain* |  |  |  |  |
| TD (s) | 16.6 ± 4.1 | 17.9 ± 4.9 | 0.575 | 0.121 |
| Baseline $\dot{V}\text{O}_2$ (L.min<sup>-1</sup>) | 0.97 ± 0.05 | 0.77 ± 0.09 | **< 0.001** | **1.516** |
| Amplitude (L.min<sup>-1</sup>) | 0.83 ± 0.19 | 0.60 ± 0.18 | **0.024** | **0.477** |
| Amplitude (ml.kg<sup>-1</sup>.min<sup>-1</sup>) | 11.0 ± 1.6 | 9.7 ± 2.6 | 0.230 | 0.323 |
| Amplitude (% $\dot{V}\text{O}_{2\text{peak}}$) | 24 ± 3 | 24 ± 5 | 0.949 | 0.018 |
| $\dot{V}\text{O}_2$ gain (ml.min<sup>-1</sup>.W<sup>-1</sup>) | 10.2 ± 0.7 | 10.5 ± 1.1 | 0.587 | 0.130 |
| $\tau$ (s) | 27.9 ± 7.5 | 24.8 ± 6.6 | 0.385 | 0.160 |
| $r^2$ | 0.965 ± 0.032 | 0.947 ± 0.034 | | |
| *Heavy Intensity Domain* |  |  |  |  |
| TD (s) | 15.0 ± 4.8 | 14.5 ± 4.5 | 0.858 | 0.033 |
| Baseline $\dot{V}\text{O}_2$ (L.min<sup>-1</sup>) | 1.00 ± 0.06 | 0.85 ± 0.11 | **0.005** | **0.894** |
| Amplitude (L.min<sup>-1</sup>) | 1.50 ± 0.38 | 0.96 ± 0.25 | **0.005** | **0.541** |
| Amplitude (ml.kg<sup>-1</sup>.min<sup>-1</sup>) | 19.9 ± 4.7 | 15.6 ± 3.7 | 0.060 | 0.351 |
| Amplitude (% $\dot{V}\text{O}_{2\text{peak}}$) | 43 ± 5 | 38 ± 7 | 0.179 | 0.355 |
| $\dot{V}\text{O}_2$ gain (ml.min<sup>-1</sup>.W<sup>-1</sup>) | 9.4 ± 0.8 | 9.3 ± 0.9 | 0.766 | 0.059 |
| $\tau$ (s) | 28.8 ± 7.9 | 27.2 ± 4.4 | 0.633 | 0.075 |
| $r^2$ | 0.961 ± 0.031 | 0.927 ± 0.030 | | |
| SC Amplitude (L.min<sup>-1</sup>) | 0.38 ± 0.16 | 0.28 ± 0.12 | 0.158 | 0.250 |
| SC Amplitude (% $\dot{V}\text{O}_{2\text{peak}}$) | 11.9 ± 6.9 | 10.8 ± 3.2 | 0.686 | 0.061 |
| SC Amplitude (% end exercise $\dot{V}\text{O}_2$) | 13.6 ± 6.5 | 12.9 ± 3.9 | 0.822 | 0.035 |

SC: slow component, $\tau$ : time constant, TD: time delay, $\dot{V}\text{O}_2$ : rate of oxygen consumption

In absolute units (L.min^-1^) males experienced greater amplitudes of V□O_2_ during the phase II kinetics (*p* ≤ 0.024), however when this was made relative to the individual (V□O_2_ gain and % V□O_2peak_) no sex differences were observed (*p* ≥ 0.179). Similarly, the V□O_2_ slow component amplitude was not different between sexes in relative units (*p* = 0.686). As visualised in Figure 1, there were no sex differences in τV□O_2_ in either the moderate (*p* = 0.385) or heavy (*p* = 0.633) intensity domains. Baseline V□O_2_ was slightly elevated at the onset of heavy intensity exercise compared to the onset of moderate intensity exercise for females (mean difference: 0.08 L.min^-1^, *p* = 0.028), whereas male baseline V□O_2_ was not different (*p* = 0.312).

**Figure 1:**
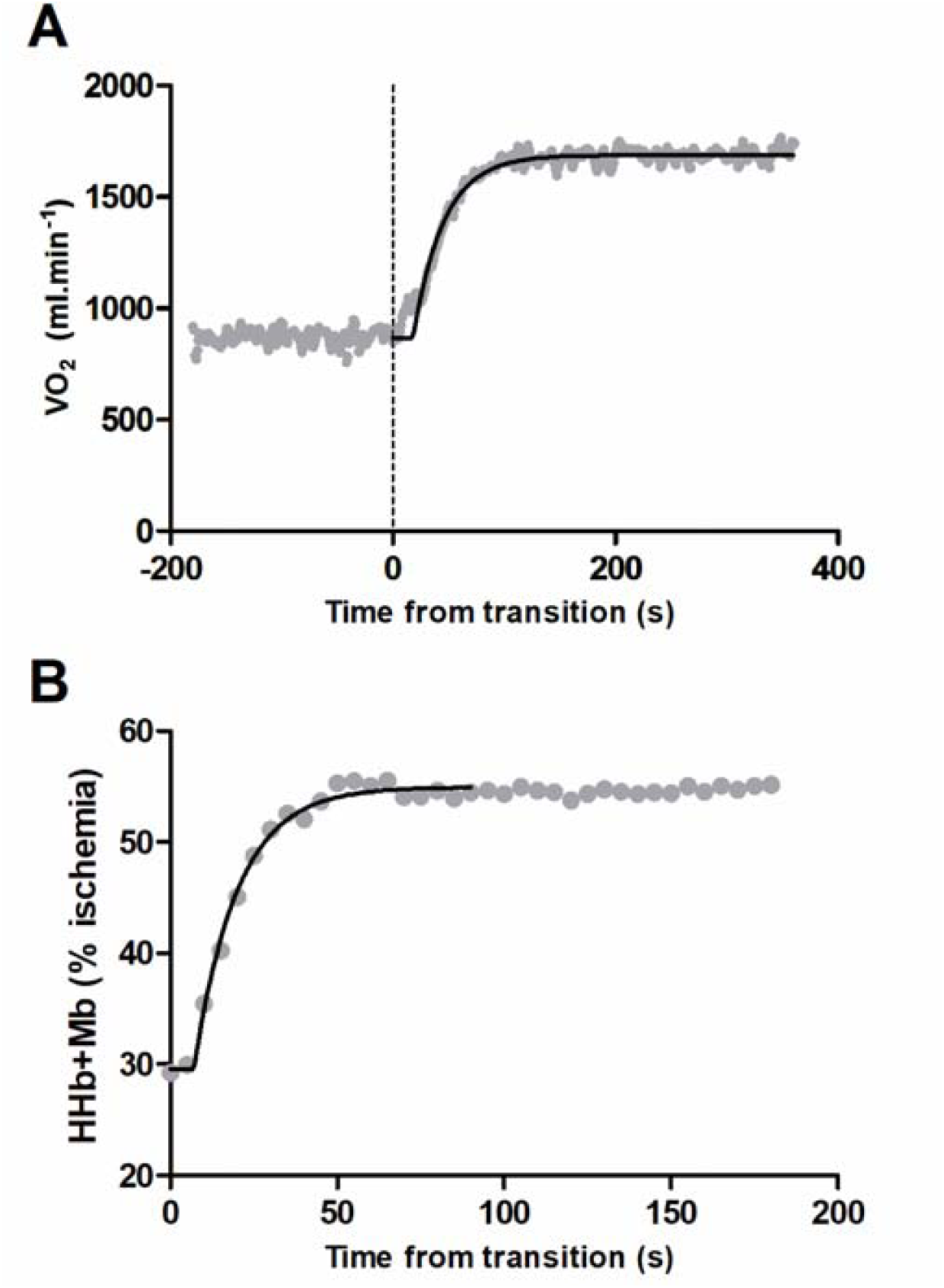
Visualisation of the monoexponential curve fitting procedures for a representative participant’s data in the moderate intensity domain. Panel A describes the V□O_2_ data (1 Hz) and Panel B describes the HHb+Mb data (0.2 Hz).

**Figure 2:**
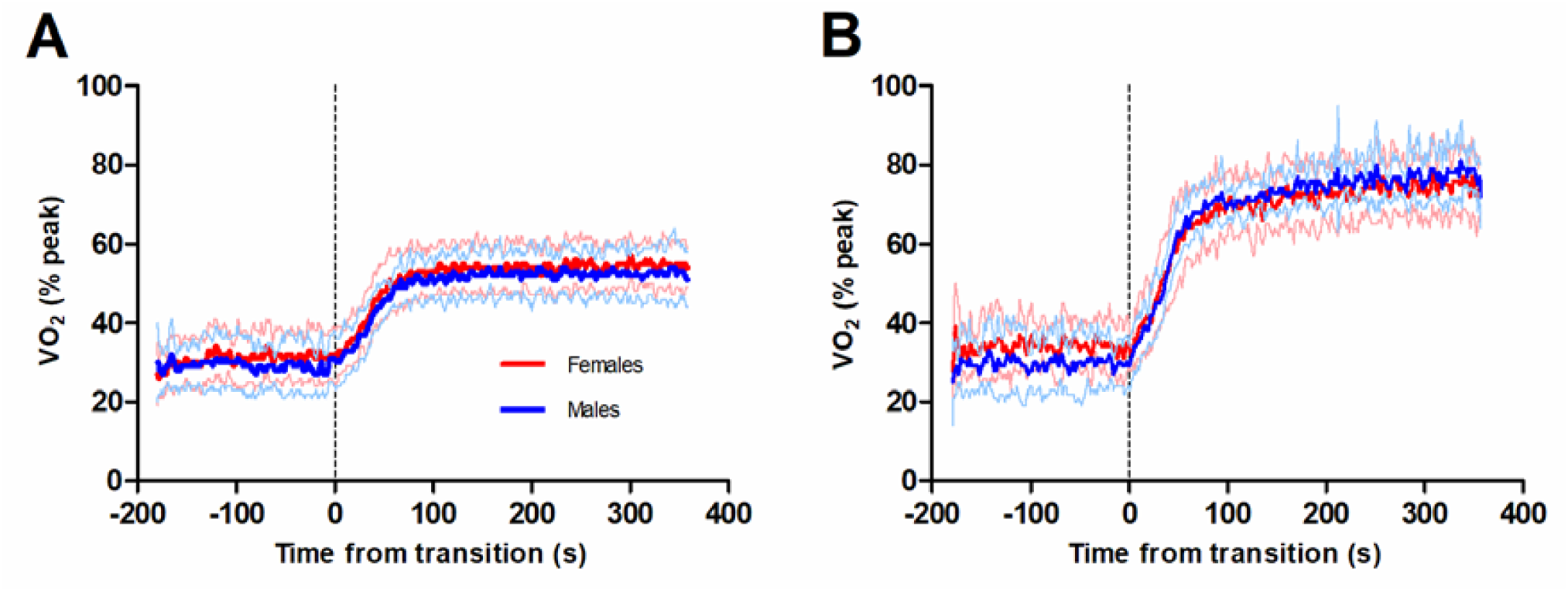
Group mean V□O_2_ data from males (blue, n=8) and females (red, n=8) during moderate (Panel A) and heavy (Panel B) intensity transitions. The bold lines represent group means and the thin lines represent standard deviation.

### Oxygen Extraction Kinetics

The transition from unloaded pedalling to moderate and heavy intensity cycling elicited an increase in HHb+Mb concentration (Figure 3), and the monoexponential curve used to describe the increase in HHb+Mb in males and females demonstrated excellent r^2^ values (Table 3). One female’s data had to be removed due to issues with the NIRS signal, resulting in n = 7 females being used for NIRS analyses.

**Figure 3:**
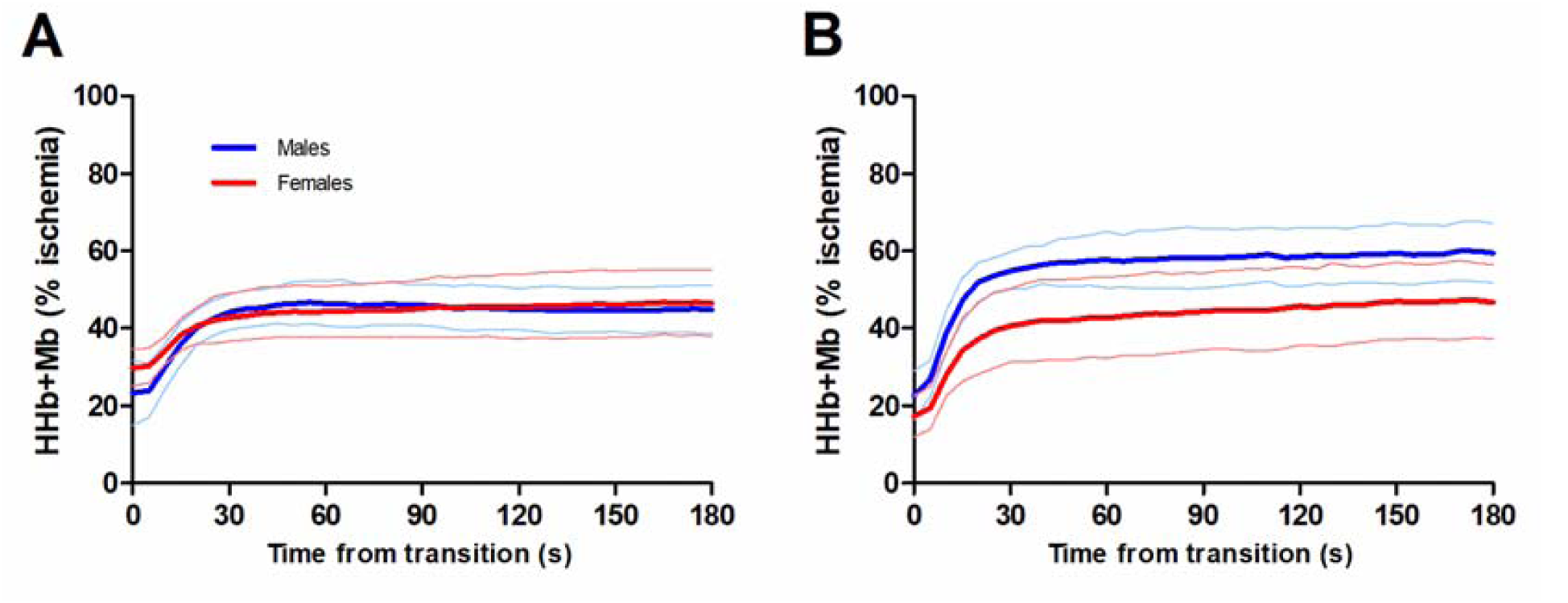
Group mean HHb+Mb data from males (blue, n=8) and females (red, n=7) during moderate (Panel A) and heavy (Panel B) intensity transitions. The bold lines represent group means and the thin lines represent standard deviation.

**Table 3:** Data from the monoexponential modelling of deoxyhaemoglobin kinetics during moderate and heavy intensity transitions.*Figure 3*: Group mean HHb+Mb data from males (blue, n=8) and females (red, n=7) during moderate (Panel A) and heavy (Panel B) intensity transitions. The bold lines represent group means and the thin lines represent standard deviation.

|  | Males (n = 8) | Females (n = 7) | P value | Cohen's d |
| --- | --- | --- | --- | --- |
| *Moderate Intensity Domain* |  |  |  |  |
| TD (s) | 8.7 ± 1.5 | 9.6 ± 2.6 | 0.447 | 0.212 |
| Baseline HHb+Mb (% ischemia) | 23 ± 8 | 29 ± 7 | 0.186 | 0.251 |
| Amplitude (% ischemia) | 21 ± 7 | 17 ± 5 | 0.225 | 0.219 |
| τ (s) | 8.1 ± 2.8 | 10.0 ± 3.8 | 0.274 | 0.260 |
| r<sup>2</sup> | 0.974 ± 0.022 | 0.956 ± 0.037 |  |  |
| *Heavy Intensity Domain* |  |  |  |  |
| TD (s) | 6.2 ± 2.8 | 7.0 ± 2.3 | 0.582 | 0.101 |
| Baseline HHb+Mb (% ischemia) | 23 ± 6 | 17 ± 7 | 0.146 | 0.332 |
| Amplitude (% ischemia) | 36 ± 9 | 30 ± 8 | 0.193 | 0.252 |
| τ (s) | 11.3 ± 3.7 | 12.2 ± 4.0 | 0.665 | 0.090 |
| r<sup>2</sup> | 0.983 ± 0.011 | 0.979 ± 0.023 |  |  |
| SC Amplitude (% ischemia) | 14 ± 5 | 16 ± 4 | 0.432 | 0.142 |

HHb+Mb: deoxyhaemoglobin and myoglobin, SC: slow component, τ: time constant, TD: time delay consumption

**Table 4:** Data from near-infrared spectroscopy before and during the moderate and heavy intensity exercise transitions in males (n = 8) and females (n = 7).

|  |  | Moderate Intensity |  | Heavy Intensity |  |  |
| --- | --- | --- | --- | --- | --- | --- |
|  |  | Unloaded Pedalling | End Stage | Unloaded Pedalling | 90 secs | End Stage |
| **HbO<sub>2</sub> + MbO<sub>2</sub>**<br>(%ischemia) | Male | 57 ± 6 | 53 ± 7 | 68 ± 11* | 54 ± 10*<sup>#</sup> | 47 ± 10<sup>#</sup> |
|  | Female | 41 ± 10 | 44 ± 9 | 51 ± 7 | 41 ± 4<sup>#</sup> | 53 ± 8 |
| **TOI**<br>(%) | Male | 69 ± 3 | 61 ± 5<sup>#</sup> | 71 ± 4 | 57 ± 6<sup>#</sup> | 50 ± 8<sup>#</sup><sup>§</sup> |
|  | Female | 75 ± 3* | 71 ± 6*<sup>#</sup> | 78 ± 4* | 70 ± 8*<sup>#</sup> | 69 ± 9*<sup>#</sup> |
| **TOI**<br>(% ischemia) | Male | 72 ± 4 | 60 ± 7<sup>#</sup> | 76 ± 6 | 53 ± 9<sup>#</sup> | 39 ± 10<sup>#</sup><sup>§</sup> |
|  | Female | 67 ± 9 | 53 ± 10<sup>#</sup> | 77 ± 6 | 52 ± 9<sup>#</sup> | 51 ± 17<sup>#</sup> |

HbO<sub>2</sub>+MbO<sub>2</sub>: oxygenated haemoglobin and myoglobin; TOI: tissue oxygenation index; \* = greater than the opposite sex ( $p < 0.05$ ); <sup>#</sup> = lower than unloaded pedalling ( $p < 0.05$ ); <sup>§</sup> = lower than 90 secs.

As shown in Figure 3, the phase II amplitude of HHb+Mb increase in both intensity domains was not different between sexes (*p* ≥ 0.193). Similarly, there was no sex difference in the amplitude of the slow component in the heavy intensity domain (*p* = 0.432). The time constant for phase II HHb+Mb kinetics (τHHb+Mb) was also not different between males and females in both intensity domains (*p* ≥ 0.274). Baseline HHb+Mb was slightly lower at the onset of heavy intensity exercise for females compared to the onset of moderate exercise (mean difference: 12% ischemia, *p* = 0.002), but not for males (*p* = 0.583).

### Near-infrared spectroscopy

During the moderate intensity cycling, a significant effect of sex was observed for HbO_2_+MbO_2_ (F_1,13_ = 10.85, *p* = 0.006), but no time (*p* = 0.804) or sex × time interaction (*p* = 0.054) effects were observed. For TOI, significant time (F_1,13_ = 48.59, *p* < 0.001) and sex (F_1,13_ = 12.57, *p* = 0.004) effects were observed, but no sex × time interaction effect (*p* = 0.095). Post-hoc tests revealed that females had greater values before (6%, *p =* 0.002) and during the stage (10%, *p* = 0.007). However, when TOI values were normalised as % ischemia, there was a main effect of time (F_1,6_ = 115.09, *p* < 0.001), but neither the main effect of sex (*p* = 0.181) or the sex × time interaction effect (*p* = 0.381) were evident.

During heavy intensity cycling, a main effect of time was observed for HbO_2_+MbO_2_ (F_2,26_ = 17.39, *p* < 0.001), as well as sex × time interaction effect (F_2,26_ = 15.67, *p* < 0.001), but no main effect of sex (*p* = 0.052). Post-hoc tests revealed that females had lower values before (-17% ischemia, *p* = 0.004) and 90 seconds after the transition (-13% ischemia, *p* = 0.005), but not at the end of the stage (*p* = 0.216). Furthermore, HbO_2_+MbO_2_ decreased in both sexes from unloaded pedalling to 90 seconds into the transition (*p* < 0.001), but for females, this returned to baseline by the end of the stage (*p* = 0.056) whereas males remained decreased (*p* = 0.002). For TOI, main effects of time (F_2,26_ = 70.21, *p* < 0.001) and a sex × time interaction effect (F_2,26_ = 15.67, *p* < 0.001) were observed, but no main effect of sex (*p* = 0.052). Post-hoc tests revealed that females had greater values than males at all timepoints (*p* ≤ 0.005). Additionally, while males demonstrated a progressive decrease in TOI at each of the three time points (*p* ≤ 0.001), females only decreased from unloaded pedalling to 90 seconds (*p* < 0.001), then no further decrease was observed at 30 minutes (*p* = 1.000). When TOI was normalised to % ischemia, a main effect of time (F_1.37,17.78_ = 49.50 *p* < 0.001) remained, however the sex (*p* = 0.266) and sex × time interaction effect (*p* = 0.112) were not observed.

## Discussion

This study aimed to compare the kinetics of V□O_2_ and HHb+Mb during moderate and heavy intensity exercise in males and females. In contrast to the hypothesis, at the onset of exercise the phase II time constants (τ) for V□O_2_ and HHb+Mb were not different between the sexes, implying that both males and females were able to increase oxidative phosphorylation at comparable rates. In absolute units, males had larger amplitude increases than females, however when normalised to the individuals’ maximum values, the rise in V□O_2_ and HHb+Mb was not different. Combined, these data demonstrate that the oxidative response to exercise is not different between sexes, which provides mechanistic insight into previously observed sex differences in the integrative response to exercise.

Previous literature investigating sex differences in the onset kinetics of oxygen transport and utilisation conflicts with the present data, with Beltrame *et al*. (2017) demonstrating quicker τV□O_2_ and τHHb+Mb in females compared to males. One potential explanation for this discrepancy could be that Beltrame *et al*. utilised a treadmill walking task, compared to cycling. In tasks where O_2_ delivery is not a limiting factor, females often outperform males. For instance, Ansdell *et al*. (2019) showed female knee-extensors had a greater relative critical torque than males during single-limb exercise. Whereas during cycling, where O_2_ delivery is a determinant of critical power (Goulding & Marwood, 2023), this metabolic threshold was not different between sexes (Ansdell *et al*., 2020a). Whilst consensus on whether O_2_ delivery does (Hughson *et al*., 2001) or does not (Grassi, 2001) limit τV□O_2_ has not been reached, it is conceivable that during tasks where O_2_ delivery and utilisation are both determinants in the metabolic response to exercise, the superior female skeletal muscle oxidative capacity (Cardinale *et al*., 2018) and vasodilatory response to exercise (Parker *et al*., 2007) is counteracted by an inferior O_2_ carrying capacity (Murphy, 2014). Within the present data, this balance manifests as a comparable τV□O_2_ in males and females, which agrees with data from do Nascimento Salvador *et al*. (2019), who demonstrated no sex difference in τV□O_2_ during a transition from unloaded pedalling to ‘very heavy’ (60% Δ) cycling exercise.

Data from incremental exercise suggests that the poorer O_2_ delivery in females results in a greater degree of O_2_ extraction to compensate (Murias *et al*., 2013). The present data contradicts this notion, as the amplitude of phase II HHb+Mb kinetics was not different between sexes (see Table 3). However, it is important to note that Murias *et al*. noted that this sex difference only occurred once incremental exercise exceeded the respiratory compensation point (i.e., the severe intensity domain), whereas the present study compared sexes in the moderate and heavy intensity domains. The lack of a sex difference in the phase II amplitude for HHb+Mb kinetics contradicts previously published NIRS data that demonstrated a smaller rise in HHb+Mb and lesser decrease in TOI in females compared to males during constant-load exercise (Ansdell *et al*., 2020a). The crucial difference in methodologies employed between the previous study and the present study is the application of a ‘physiological calibration’ to negate the influence of adipose tissue thickness on NIRS signals (Ryan *et al*., 2012). Previously, the sex difference in the rise in HHb+Mb was suggested to reflect a lower oxygen cost of muscle contraction in female knee-extensors, however the present data, with more rigorous methodologies employed, refutes this. One aspect of the modelling that did demonstrate a sex difference was the reduction in HHb+Mb (and concomitant increase in V□O_2_) baseline for the heavy intensity transition in females, but not males. This could reflect a sex difference in how O_2_ utilisation is altered by prolonged or intermittent exercise (i.e., the preceding three bouts of moderate intensity cycling); however, the present study was not configured in a manner appropriate to answer that research question. Females did demonstrate lower HbO_2_+MbO_2_ levels during heavy intensity cycling, perhaps indicating a lesser O_2_ availability. This would fit with the notion that O_2_ carrying capacity is inferior in females during high-intensity exercise (Diaz-Canestro *et al*., 2022); however, the lack of difference in the speed and amplitude of HHb+Mb onset kinetics (and pulmonary V□O_2_) implies that oxygen extraction is not negatively affected by this in the moderate and heavy intensity domains.

Although not measured in the present study, the sex difference in muscle fibre type, whereby females demonstrate a greater proportional area of type I fibres (Staron *et al*., 2000; Roepstorff *et al*., 2006), appears to have not influenced either V□O_2_ or HHb+Mb onset kinetics in the present study. This would concur with data from Barstow *et al*. (1996), who demonstrated no relationship between type I fibre percentage of the *vastus lateralis* and the time constant of phase II V□O_2_ kinetics. In contrast, Pringle *et al*. (2003) observed a negative correlation between type I fibre percentage and the phase II time constant in the heavy intensity domain only. Of note is that Pringle *et al*. included a wide range of participants with ∼27 – 85% type I fibres, and when groups were split into discrete groups of low and high fibre type percentages (mean difference: 25%), the high percentage group had faster phase II kinetics. The present study was not able to quantify the sex difference in muscle fibre typology, however, previous literature has observed a 5-13% difference in type I fibre percentage of the *vastus lateralis* (Simoneau & Bouchard, 1989; Staron *et al*., 2000; Roepstorff *et al*., 2006). Therefore, it could be the case that the sex difference in muscle fibre typology is not large enough to affect the phase II V□O_2_ or HHb+Mb kinetics. Indeed, recent evidence suggests that during exercise normalised to metabolic thresholds, sex differences in muscle fibre typology do not influence fatigability (McDougall *et al*., 2023).

Muscle fibre typology has previously been demonstrated to affect the amplitude of the slow component within the heavy intensity domain, as individuals with a lower type I fibre percentage experience larger rises in V□O_2_ during constant-load exercise (Barstow *et al*., 1996; Pringle *et al*., 2003). It is suggested that the slow component is mechanistically underpinned by factors such as additional motor unit recruitment (Poole *et al*., 1994; Burnley *et al*., 2002) to compensate for fatigue-related changes in muscle metabolism. For instance, muscle PCr stores demonstrate a similar slow component in depletion during heavy intensity exercise (Rossiter *et al*., 2002). Given that female knee-extensors appears more fatigue-resistant and demonstrate lesser rises in the amplitude of surface electromyography during constant-load exercise (Ansdell *et al*., 2017; Ansdell *et al*., 2019; Ansdell *et al*., 2020a), we hypothesised that the relative amplitude of the slow component would be greater in males to reflect a greater rate of metabolic disturbance. However, as is evident in Tables 2 and 3, no sex difference was observed in the relative slow component amplitude, implying that there was no difference in the metabolic response to constant-load exercise.

The lack of sex differences in either the phase II kinetics or slow component amplitude collectively suggest that the oxidative response to exercise was not different between males and females. Data on this topic is sparse, and due to the nature of methods such as magnetic resonance spectroscopy (MRS), limited to single-joint, isometric muscle contractions. Previous literature using this technique to study muscle metabolic changes during a 60 s contraction of the dorsiflexors showed no sex difference in changes in PCr, Pi, or pH (Russ *et al*., 2005). Data from muscle biopsies of the *vastus lateralis* taken before and after repeated 30s cycling sprints suggested a greater preservation of ATP concentrations in females across a ∼60 minute protocol (Esbjörnsson-Liljedahl *et al*., 2002); however the authors suggested that this was likely a result of sex differences in the 20 minute recovery periods, rather that metabolic differences during exercise. Accordingly, the same group observed no sex differences in the metabolic response to a single 30-second cycling sprint (Esbjörnsson-Liljedahl *et al*., 1999). Collectively, across multiple tasks and methodologies (MRS, biopsy and V□O_2_ kinetics), the data suggest that there is no sex difference in the bioenergetic response to high-intensity exercise. This information provides mechanistic insight into the sex differences in the integrative response to exercise. For instance, sex differences in fatigability have partly been attributed to a lesser accumulation of fatiguing metabolites (Hunter, 2014; Ansdell *et al*., 2020b). It is perhaps more accurate to suggest that previously observed sex differences in fatigue during intensity-matched exercise (Ansdell *et al*., 2020a; Azevedo *et al*., 2021) are more likely due to a greater fatigue-resistance of female muscle contractile apparatus, which experience similar degrees of metabolic stress as males. It is established that males and females differ in contractile properties such as calcium (Ca^2+^) kinetics of the sarcoplasmic reticulum (Harmer *et al*., 2014), with lower Ca^2+^ATPase activity thought to permit a more fatigue-resistant skeletal muscle profile during equivalent exercise tasks (Hunter, 2014). Therefore, the present study advances the contemporary understanding of sex differences in the integrative response to exercise and provides mechanistic insight into previously observed phenomena.

These data have applications across the spectrum of health and disease. For example, those prescribing steady state exercise to improve skeletal muscle performance in athletes or patients might not need to account for the sex of their participants (Gloeckl *et al*., 2022; Furrer *et al*., 2023). This statement should however, be caveated by the fact that evidence regarding the influence of sex on long-term adaptation to exercise is sparse (Ansdell *et al*., 2020b). Indeed, one area for further exploration is the bioenergetic response to exercise within the severe intensity domain, where sex differences in fatigability have previously been observed (Ansdell *et al*., 2020a; Azevedo *et al*., 2021). The employment of complementary techniques to quantify O_2_ delivery (for instance, the present study did not quantify total Hb+Mb) and muscle fibre typology could also provide greater insight into the influence of sex on the O_2_ cascade in a variety of tasks.

Previous evidence from pre-pubertal children and adolescents suggests that boys have a lower τV□O_2_ and V□O_2_ slow component than girls (Armstrong & Barker, 2009), which contradicts the present findings in adults. Previous studies comparing children and adults also found that intramuscular PCr kinetics were similar in males and females regardless of age (Willcocks *et al*., 2010), implying similar oxidative capacity. Interestingly, the same study found a greater ‘PCr cost’ (mM.W^-1^) in females compared to males suggesting a greater inefficiency in females, which contradicts the present findings. Caution is urged when comparing data from Willcocks *et al*. (2010) and the present study, however, given the nature of exercise (single joint vs. whole body), the lack of matching for aerobic fitness, and the low sample size (6 males vs. 5 females).

Finally, hormonal status was not an exclusion criterion or controlled for within female participants in the present study. We based this decision on evidence from Mattu *et al*. (2020) who demonstrated that V□O_2_ kinetics did not differ between the follicular and luteal phases of the eumenorrheic menstrual cycle, or with oral contraceptive usage. However, we do acknowledge that the aforementioned study only investigated the moderate intensity domain, and therefore might not apply to V□O_2_ kinetics in the heavy intensity domain. In the heavy intensity domain, contributing factors to the V□O_2_ slow component (e.g., substrate utilisation) might be affected by hormonal status, although the available evidence is conflicting (Oosthuyse *et al*., 2023). Therefore, further research is required to determine whether endogenous and exogenous hormones influence the oxidative response to high-intensity exercise.

## Conclusion

The present study aimed to compare the oxygen extraction and uptake kinetics during moderate and heavy intensity cycling exercise. Contrary to our hypotheses, no sex differences were observed in either the phase II or slow component kinetics for V□O_2_ or HHb+Mb. The lack of sex difference implies that males and females do not experience different oxidative responses to exercise, which provides mechanistic insight into previously observed phenomena such as the sex difference in fatigability. Furthermore, based on these data and others demonstrating no hormonal influences (Mattu *et al*., 2020), we suggest that there is no rationale for the exclusion of female participants in research investigating cardiopulmonary responses to exercise.

## Acknowledgements

The authors wish to thank the participants for their time and effort, as well as the technical staff within the Department of Sport, Exercise and Rehabilitation for their support.

## Funding

This project was supported by a Physiological Society Research Springboard Studentship awarded to MSP, as well as Erasmus+ funding awarded to LB (2020-1-DE01-KA103-005569). PA is supported by the UK Office for Veteran’s Affairs (G2-SCH-2022-11-12245).

## Disclosures

There are no conflicts of interest, financial or otherwise.

## Author Contributions

MSP, ZW, MB, and PA conceived and designed the research. MSP, LB, EH, LH, and PA performed the experiments. LB, EH, LH, ZW, and PA analysed data. MSP, LB, EH, LH, ZW, MB, and PA interpreted results of the experiments. PA drafted the manuscript. MSP, LB, EH, LH, ZW, MB, and PA edited and revised the manuscript. All authors approved the final version of the manuscript.

## Notes

### Competing Interest Statement

The authors have declared no competing interest.

### Summary of Updates

The results section was updated to include data from near-infrared spectroscopy regarding muscle oxygenation status at key time points before and during the square wave exercise transitions, as well as some additional normalisation of VO2 data. Other minor amendments to the manuscript are found throughout.

